# Feasibility of Functional MRI at Ultralow Magnetic Field via Changes in Cerebral Blood Volume

**DOI:** 10.1101/366955

**Authors:** Kai Buckenmaier, Anders Pedersen, Paul SanGiorgio, Klaus Scheffler, John Clarke, Ben Inglis

**Author notes:** High-Field Magnetic Resonance Center, Max Planck Institute for Biological Cybernetics, Tübingen, Germany (Kai Buckenmaier). Illumina, Inc., 5200 Illumia Way, San Diego CA 92122. Declaration of interest:* JC is the holder of US patent 7466132, “SQUID-detected NMR and MRI at ultralow fields.”.

## Abstract

We investigate the feasibility of performing functional MRI (fMRI) at ultralow field (ULF) with a Superconducting QUantum Interference Device (SQUID), as used for detecting magnetoencephalography (MEG) signals from the human head. While there is negligible magnetic susceptibility variation to produce blood oxygenation level-dependent (BOLD) contrast at ULF, changes in cerebral blood volume (CBV) may be a sensitive mechanism for fMRI given the five-fold spread in spin-lattice relaxation time (*T*_1_) values across the constituents of the human brain. We undertook simulations of functional signal strength for a simplified brain model involving activation of a primary cortical region in a manner consistent with a blocked task experiment. Our simulations involve measured values of *T*_1_ at ULF and experimental parameters for the performance of an upgraded ULFMRI scanner. Under ideal experimental conditions we predict a functional signal-to-noise ratio of between 3.1 and 7.1 for an imaging time of 30 minutes, or between 1.5 and 3.5 for a blocked task experiment lasting 7.5 minutes. Our simulations suggest it may be feasible to perform fMRI using a ULFMRI system designed to perform MRI and MEG *in situ.*

## Introduction

The goal of functional neuroimaging is to detect and localize brain activity. Ensemble neuronal activity generates cellular ionic currents that can be measured on the surface of the head using electroencephalograpy (EEG) or close to the scalp using magnetoencephalography (MEG), each with a temporal precision of milliseconds (da Silva, 2013). As imaging modalities, however, EEG and MEG are ill posed since their sources cannot be localized precisely from measurements of their fields at a distance (Cohen, 2017). Recent technical developments have shown that it is possible to perform MEG and magnetic resonance imaging (MRI) with an integrated apparatus, using the same set of Superconducting QUantum Interference Device (SQUID) gradiometers for both signals (Volegov et al., 2004; Zotev et al., 2008; Vesanen et al., 2013). Localization with MRI is well defined. The MR images are obtained at an ultra-low magnetic field of tens to hundreds of microtesla, comparable to Earth’s magnetic field. To date, all ultralow field (ULF) MRI of the human brain has been anatomical (Volegov et al., 2004; Zotev et al., 2008; Vesanen et al., 2013; Zotev et al., 2009; Inglis et al., 2013). It has been suggested that the weak fluctuating magnetic fields observed with MEG might be detected directly with ULFMRI from the tiny frequency shift in the signal (Neuronal Current Imaging, NCI) (Kraus et al., 2008; Cassara et al., 2008; Cassara et al., 2009; Körber et al., 2013). Although recent advances in SQUID technology have reduced the noise closer to the predicted required level (Körber et al., 2016), so far there has been no demonstration of NCI in a human brain.

Brain activity can also be estimated from changes in metabolism, which itself can be approximated by alterations in the regional blood supply to part of the brain (Buxton, 2002). Other than the systematic anatomical displacement of the vasculature from the neuronal populations of interest, vascular MRI signals can be located with millimeter precision in human functional MRI (fMRI) experiments, but temporal resolution is constrained by sluggish hemodynamics that are delayed by several seconds following a short stimulus (Goense et al., 2016; Huber et al., 2017). The standard approach to fMRI at high magnetic fields uses the blood oxygenation level-dependent (BOLD) contrast mechanism, which relies on a change in magnetic susceptibility of the venous blood draining a site of neural activity (Ogawa et al., 1990; Kwong et al., 1992, Bandettini et al., 1992; Ogawa et al., 1992). The magnitude of the BOLD effect scales at least linearly with the external magnetic field strength and is thus predicted to be vanishingly small at ULF (van der Zwaag et al., 2009). Negligible BOLD at ULF suggests other vascular approaches could be adopted without concomitant

BOLD signals to complicate interpretation. One might use arterial spin labeling (ASL) to perform fMRI at ULF, using cerebral blood flow (CBF) changes (Kim, 1995; Wong et al., 1997). Given, however, the longitudinal relaxation time *T*_1_ of blood, approximately 200 ms at 130 μT (Inglis et al., 2013), and a typical bolus arrival time from arteries to capillaries of more than 500 ms, we predict that ASL using spatial labeling will have a poor signal-to-noise ratio (SNR) at ULF. Similarly, while it is possible in principle to label blood water arbitrarily close to brain tissue using velocity-selective labeling for ASL (Wong et al., 2006), the sensitivity also decreases as blood *T*_1_ decreases. The ASL-based approaches benefit from higher magnetic fields, where blood *T*_1_ is longer.

An alternative hemodynamic response is the change in the CBV concomitant with modulation of CBF (Grubb et al., 1974; Chen and Pike, 2009). In CBV-based functional imaging the signal changes arise from the difference in *T*_1_ between brain tissue, cerebrospinal fluid (CSF) and blood. The CBV response can be measured with the vascular space occupancy (VASO) method (Lu et al., 2003), aimed at nulling the blood compartment selectively, or by the closely related scheme using gray matter nulling (GMN) (Wu et al., 2008; Shen et al., 2009; Shen et al., 2012). Our group has determined a *T*_1_-difference between CSF and blood of more than a factor of five, and between blood and brain tissue of more than a factor of two (Inglis et al., 2013), making CBV-based sequences an attractive possibility for ULF-fMRI.

A vascular approach to functional ULFMRI might be interleaved with multichannel-MEG detection to take advantage of the different temporal dynamics of the neuronal and vascular responses. A combined MEG and ULFMRI system might permit direct detection of neural events with MEG, followed a few seconds later by the vascular signal changes for detection via functional MRI. The combination of MEG and functional ULFMRI could provide simultaneously fast sampling rate and well defined localization for assessing brain activity non-invasively. We were therefore motivated to explore methods for performing functional MRI at ULF, and to estimate the magnitude of signal changes that would be required to establish functional localization. Here we present simulations of an idealized experiment using CBV-based functional contrast with a simple brain activation paradigm.

## Materials and Methods

### ULFMRI Instrument

Our ULFMRI instrument (McDermott et al., 2004; Clarke et al., 2007; Inglis et al., 2013) is surrounded by a 1.5-mm thick cubical aluminum shield, 1.8 m on a side, with the seated subject’s head at the center [Fig. 1A, B]. Magnetic fields are generated by copper coils. Two pairs of coils (*B_Cx_, B_Cy_*) wound on the faces of the cube cancel the Earth’s field over the imaging region in the *x*- and *y*-directions, and a Helmholtz pair reinforces the *z*-component of the Earth’s field to produce the imaging field *B*_0_, usually about 130 μT. A Maxwell pair produces the gradient field *G_z_* ≡ *∂B_z_/∂z,* and two sets of planar gradient coils produce *G_x_ ≡ ∂B_z_/∂x* and *G_y_ ≡ ∂B_z_/∂y.* The gradient fields are typically 100 μT/m. An excitation coil provides oscillating pulses *B*_1_ along the *y*-axis to manipulate the spins. To magnetize the protons, a water-cooled coil generates a prepolarization field *B_p_,* approximately 80 mT at the subject’s head, along the *x*-axis. After the polarizing field is switched off adiabatically in 10 ms, an imaging pulse sequence encodes the magnetization (Inglis et al., 2013). The nuclear magnetic resonance (NMR) signal is detected by a superconducting, second-derivative axial gradiometer [Fig. 1C] coupled to a dc SQUID (see Supplementary material) immersed in liquid He at 4.2 K in a fiberglass dewar placed directly above the head [Fig. 1B]. The radius of the gradiometer loops is *R* = 38 mm and the separation of the lowest and uppermost loops 150 mm. The total ambient and intrinsic magnetic field noise of the detector, referred to the lowest loop, is typically 0.7 fTHz^−1/2^.

**Fig. 1.**
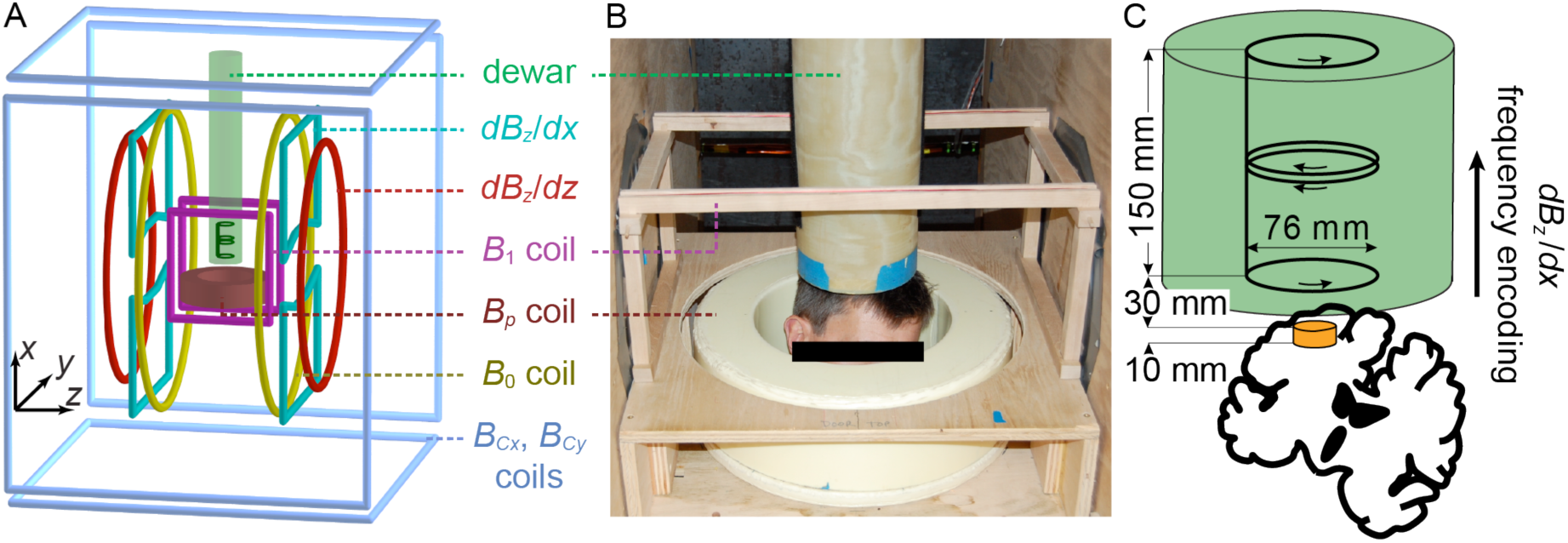
(A) Schematic and (B) photograph of the ULF-MRI system. (C) Schematic cross-section of a brain with the second-order gradiometer in a fiberglass dewar (green cylinder) situated above. The orange disk indicates the ROI, comprising the hand region of primary motor cortex.

### Model of Activated Tissue

As a representative fMRI experiment we consider stimulation of the hand region of the primary motor cortex (PMC), for example cued blocks of a finger-tapping task. It is convenient to use a manual task in our ULFMRI because the primary motor cortices are located toward the top of the brain, along the precentral gyrus, and a small head tilt locates a region-of-interest (ROI) directly under the sensor [Fig 1B,C]. The separation of the left and right motor cortices across the brain hemispheres, connected via the corpus callosum, makes left-hand finger tapping a robust control for right-hand finger tapping, or *vice versa,* while for a simple motor sequence the degree of body and head motion is likely to be reasonably consistent between the two conditions.

Since we are interested only in PMC, we constrain the ROI to reside within a 10-mm thick band defined by frequency encoding parallel to the axis of the gradiometer [Fig. 1C]. The distance between the top of PMC and the lowest loop of the gradiometer is 30 mm, that is, 10 mm from the loop to the scalp and an assumed distance of 20 mm from the scalp to the top of PMC. We choose the head orientation to provide the highest possible signal from PMC [orange disk, Fig 1C]. With active tissue in PMC restricted to a band *L* = 10 mm thick we can model the ROI as a cylinder with height *L* and radius *R*_ROI_ = 10 mm, which is a reasonable approximation of the extent of hand activation patterns observed with conventional BOLD-based fMRI (Budde et al., 2014). The assumed 3D localization of an ROI from the interaction of a frequency encoding gradient with the receive profile of the gradiometer is suitable for the calculation of functional signal change, but implies that there is no movement between the baseline and activation states. All signal outside the ROI is thus assumed to cancel perfectly.

The CBV-based fMRI methods rely on small changes from a resting CBV of 4-6% in gray matter (GM) (Buxton, 2002, p.427). A local increase in CBV causes displacement of water from adjacent extracellular fluid, a compartment that is assumed to be in direct contact with CSF in sulci (Krieger et al., 2012; Scouten and Constable, 2008). Thus, for relatively low spatial resolution, the *T*_1_ of the CSF fraction must be taken into account, in addition to the difference in *T*_1_ of blood and tissue (Scouten and Constable, 2008; Lu et al., 2013). The CSF is well approximated by pure water, with a spin density we define as unity. Blood is approximately 87% water by volume (Lu et al., 2002), while GM and WM have spin densities of about 0.89 and 0.73, respectively (Simpson et al., 2001, p. 89, Shen et al., 2012). We assume that the stated water proton densities represent freely diffusing, NMR-visible water compartments. The *T*_1_ values are adapted from our prior work (Inglis et al., 2013). We assume *B*_0_ = 130 μT, as used previously. For *T*_1_ in the polarizing field we modify values measured at 80 mT according to the power law dependencies reported by Rooney *et al.* (Rooney et al., 2007), for *B_p_* = 160 mT (as explained below). Table 1 summarizes the relative spin densities and *T*_1_ values. We assume that GM and WM have identical *T*_1_-values at ULF.

**Table 1.** Tissue properties for the simplified brain activation model.

| Tissue Type | $T_1 (B_0 = 130 \mu\text{T})$ | $T_1 (B_p = 160 \text{ mT})$ | Relative Spin Density |
| --- | --- | --- | --- |
| CSF | 1770 ms | 4360 ms | 1.00 |
| Blood | 190 ms | 570 ms | 0.87 |
| Gray Matter | 85 ms | 589 ms | 0.89 |
| White Matter | 85 ms | 589 ms | 0.73 |

Properties of tissue types for model of human brain. Spin-lattice relaxation times $T_1$ at $B_0 = 130 \mu\text{T}$ and $B_p = 160 \text{ mT}$ , and spin densities relative to water. Values for $T_1(B_p)$ were scaled by 1.267, 1.3 and 1.3 for blood, GM and WM, respectively, compared to measured values reported at 80 mT in Ref. (Inglis et al., 2013), based on the power law dependencies found by Rooney *et al.* (Rooney et al., 2007). The $T_1$ for CSF has negligible $B_p$ dependence from 80 to 160 mT.

The signal change on activation depends on the volume fraction of activated tissue—in this case the hand region of PMC—relative to all other tissues generating signal in the ROI. CSF may be present in sulci or as interstitial fluid distributed in GM, neither of which are resolved in our 3142 mm^2^ ROI. Water in these compartments is assumed to exchange freely. We also note that while the CSF fraction may not be an important consideration for VASO performed with high spatial resolution at high magnetic field, *T*_1_ for CSF is an order of magnitude longer than *T*_1_ of other brain components at ULF, and could produce a large relative signal here. We thus take a grand average volume fraction of 0.10 for CSF. For blood, we consider both a microvascular compartment that can be manipulated via CBV changes, producing concomitant volume changes in other tissue compartments, together with a passive fraction occupied by large caliber vessels. It is generally accepted that microvascular blood has a volume fraction of around 0.05 in GM (Krieger et al., 2012). We assume a further volume fraction of 0.05 is occupied by blood in large caliber arteries and veins that are included unintentionally in our 3142 mm^2^ ROI. During activation, we assume that the microvascular blood volume fraction increases by 20% (Krieger et al., 2012; Lu et al., 2013; Huber et al., 2015), from 0.05 to 0.06, while the large caliber volume fraction is unchanged at 0.05, resulting in an increase from (0.05+0.05) = 0.10 at baseline to (0.06+0.05) = 0.11 total blood volume fraction on activation.

In methods such as VASO and GMN it is usually assumed that an increase of CBV is offset by a concomitant decrease in the volume occupied by GM tissue. It has been suggested, however, that the CSF volume fraction may also change to accommodate the CBV increase in some brain regions (Scouten and Constable, 2008). Since GM and CSF have widely different *T*_1_-values at ULF, we simulate two extreme possibilities: (i) the volume fraction of CSF changes from 0.10 to 0.09 and the tissue volume fraction is unchanged at 0.80, and (ii) the tissue volume fraction changes from 0.80 to 0.79 and the CSF compartment is unchanged at 0.10. In this manner, we can simulate the extreme signal changes that would accompany a 20% increase in CBV without needing to determine the actual redistribution of tissue water.

With BOLD-based fMRI at high magnetic fields, the weak functional signals are usually evaluated using a measurement time series on which one performs statistical tests. This is a form of signal averaging. At ULF, we also expect a small difference between the signal upon activation and the baseline signal level. We calculate a signal change upon activation for a spatially localized signal and the time required to obtain that signal. Then, for simplicity, we assume conventional time averaging of pairs of activation and baseline signal acquisitions to increase the functional contrast-to-noise ratio.

### Sensitivity and Functional Contrast-to-Noise Ratio

We first consider the sensitivity of our detector to sample magnetization. We calculate the signal flux through a loop using Φ(*t*) = ∫_RQI_ ***β***(***r***) · ***M***(***r****, t*)*dV* (Haacke et al., 1999). Here, ***M***(***r****, t*) is the magnetization of the sample and ***β***(***r***) is the magnetic field per unit current that would be produced by the coil at point ***r*** which, by reciprocity, is equivalent to the receive field for the loop. Given the relatively small dimensions of the ROI and its distance from the lower loop of the gradiometer, we assume a constant receive field over the ROI. The assumption of homogeneous magnetization over the ROI and a homogeneous receive field reduces the problem to a loop that detects the magnetic flux of an equivalent magnetic dipole. This is analogous to the inverse of a simple MEG experiment in which we know the dipole strength and location but seek the field detected in the pickup loop. The strength of the dipole is given by the volume and average spin density of our ROI and by the magnetization produced by the pulse sequence used for functional contrast, as discussed below.

The maximum magnetization ***M***(***r****, t*) = *M*_echo_ is measured at the top of a spin echo. The signal intensity is proportional to *M*_echo_. Furthermore, since the magnetization at the echo top is perpendicular to the precession field, only the receive field profile *β*_⊥_(**r**) perpendicular to the precession field is relevant. Dividing the signal flux Φ(*t*) by the pickup loop area *A_p_*, we find the detected magnetic field strength

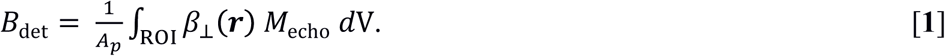

We calculate the receive field *β*_⊥_ (***r***) *= β*_⊥_ using the Biot-Savart law, with *β*_⊥_ *= μ*_0_*R*^2^/2(*R*^2^ *+ d*^2^)^3/2^, where *μ*_0_ is the vacuum permeability, and *d* is the distance between the centers of the ROI and the lowest gradiometer loop. The integral itself is the volume of the ROI, 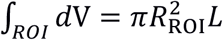. The detected magnetic field strength is

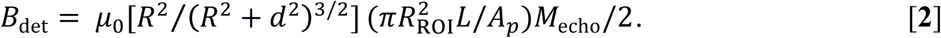

We neglect the magnetic fields coupled into the upper three loops of the gradiometer, leading to an overestimate of the signal magnitude no larger than 15%.

We next calculate the magnetization *M*_echo_ of the ROI at the time of signal detection. For functional ULFMRI the prepolarizing pulse sequence is followed by a frequency-encoded spin echo sequence [Fig. 2]. There is a variable waiting time *T_w_* between the start of the *B_p_* ramp-down and the excitation *B*_1_ pulse. A time *T_w_* of 10 ms allows an adiabatic sweep of the field direction from along *B_p_*(*x*) to along *B*_0_(*z*), and relay switching in the polarizing coil circuit. Because of experimental limitations—primarily the duration of *B*_1_ fields and the necessity of using low amplitude imaging gradients to minimize concomitant field effects (Myers et al., 2005)—the system has a minimum echo time, *TE,* of 20 ms. During *T_w_* and *TE* the signal decays as a function of *T*_1_ and *T*_2_, respectively. We assume that *T*_1_ ≈ *T*_2_ for all components of the head at 130 μT, as observed previously (Inglis et al., 2013). For simplicity, we assume signal detection occurs for the 10 ms before and 30 ms after the echo top (*T*_acq_ = 40 ms), and that a single spin echo signal is acquired. The effective magnetic field during *T_w_* is *B*_0_.

**Fig. 2.**
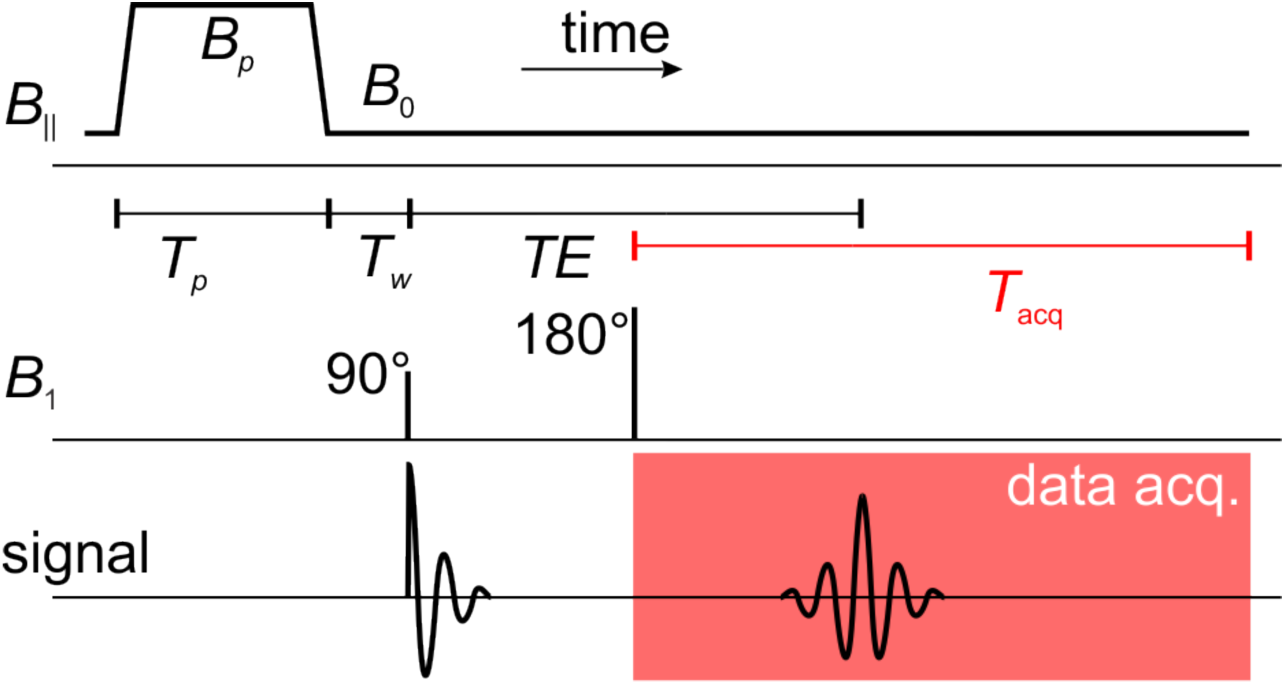
Pulse sequence for functional ULFMRI experiments. Not shown is the frequency encoding gradient *G_x_*, applied during the entire sequence. The sequence repetition time *TR* = (*T_P_* + *T_w_* + *TE*/2 + *T*_acq_). The variable delays *T_p_* and *T_w_* are swept from 0 to 10 seconds while *TE* and *T*_acq_ are kept constant.

Since at the imaging frequencies the magnetic field noise spectrum 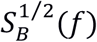 of our detector is frequency-independent, the root mean square (rms) magnetic field noise is 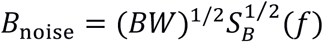 (Clarke and Braginski, 2004). Here, the acquired signal bandwidth is *BW* = *γG*_freq_*L*/2*π*, where *γ*/2*π* the proton gyromagnetic ratio, *G*_freq_ is the strength of the frequency encoding gradient and *L* is the height of the cylindrical ROI. We define a functional contrast-to-noise ratio

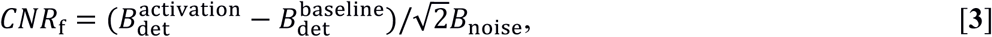

where 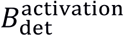 and 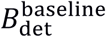 are the activation and baseline signal levels calculated from Eq. **2**. The factor √2 accounts for the subtraction of the two signals with uncorrelated noise. The values of *M*_echo_ used in Eq. **2** for the activation and baseline states are functions of the pulse sequence used to generate CBV-weighted contrast, and are defined below. Note that the numerator on the right side of Eq. **3** may be positive or negative, indicating that the assignment of activation and baseline states is arbitrary and that, as with conventional CBV and BOLD-based fMRI, deactivation is permissible.

### Preparation Pulse Sequences for Functional Contrast

The initial goal of our simulations was to determine the pulse sequence that maximizes *CNR*_f_. While inversion recovery (IR) is the usual approach to attaining CBV-weighted functional contrast at high static magnetic field with the VASO and GMN methods, in the case of field-cycled MRI we have greater flexibility and can optimize with respect to two (or more) different *T*_1_ values for each tissue component. The increased flexibility suggests that we should consider other sequences generating *T*_1_ contrast, namely one, two or three separate polarizing pulses with variable low-field evolution delays between them. The two- and three-pulse variants are equivalent to IR and double IR (DIR), respectively. The IR and DIR results converged with the single pulse sequence timing, however, as discussed in the Supplementary material. Here we present the single prepolarizing sequence only.

Using the Bloch equations, we calculate the evolution of the magnetization parallel to *B*_0_ for the single prepolarizing pulse sequence [Fig. 2] as

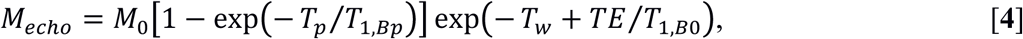

where *T*_1_,*_Bp_* and *T*_1_,*_B_*_0_ are the longitudinal relaxation times at fields *B_p_* and *B*_0_, respectively, and *M*_0_ = *ρ_s_γ*^2^*ħ*^2^*B_p_*/4*k_B_T* is the initial magnetization produced by the prepolarizing field *B_p_.* Here, *ρ_s_* is the proton density, *h* = 2π*ħ* the Planck constant, *k*_B_ the Boltzmann constant and *T* the absolute temperature. Given that the size of our existing polarizing coil is much larger than the ROI, *B_p_* varies by only about 1 mT across the entire ROI, leading us to set *B_p_* = 160 mT. We also ignore the magnetization produced by *B*_0_ = 130 μT since it is three orders of magnitude smaller than *B_p_*.

### Numerical Simulations of fMRI at ULF

We determined *CNR*_f_ for a 10-mm-long cylindrical ROI with radius 10 mm, which we assume to encompass completely the hand region of PMC. We further assume a homogeneous distribution of tissue types across the ROI, with all elements composed of grand average volume fractions of 0.8 for tissue, 0.1 for CSF and 0.1 for blood in the resting state, as explained previously, since we do not know the precise location of each fraction. We assume a body temperature of 37°C (310 K), and reference the proton densities for the brain constituents to the proton density of pure water at body temperature, 6.65×10^28^ *m*^−3^. The values *G*_freq_ = 90 μT/m and *L* = 10 mm yield *BW* = 38 Hz. As discussed in the Supplementary material, for the purposes of our simulations we assume that a next-generation ULFMRI instrument will achieve a sensor noise of 0.1 fTHz^−1/2^ and a polarizing field *B_p_* = 160 mT in the ROI.

We performed simulations using a custom routine written in Matlab (MathWorks, Natick, MA). For simplicity, we first calculated *CNR*_f_ (Eq. **3**) for a single pair of activation/control measurements while sweeping the polarizing pulse durations and evolution delays from 0 to 10 s in steps of 10 ms. As with conventional MRI, the parameters yielding the maximum SNR per unit time may be different from those yielding the largest single-shot SNR, and we expect to use multiple averages to acquire a measurable functional signal. Thus, we sought the sequence that provides the maximum *CNR*_f_ per unit time. The total signal scales as √*n*, where *n* is the number of pairwise averages; the *CNR*_f_ scales with the square root of the total acquisition time for the series. We assume a total acquisition time of 30 minutes, which we have found to be tolerated by subjects sitting in our system. Hence, the final simulation results show the *CNR*_f_ obtained after 30 minutes of averaging, *CNR*_f_(30 min), taking into account the variable repetition time *TR* of the pulse sequence used for functional contrast.

With respect to signal stability, we assume there is no motion between each pair of activation and baseline measurements so that signals from all regions outside the ROI cancel perfectly after subtraction. We can thus neglect extraneous signals. Furthermore, considering only signals originating in the ROI, we obtain the largest true activation signal according to Eq. **3**. We made a further simplification concerning the temporal nature of functional signal changes in a time series. It is well known that the hemodynamic response to a brief stimulus causes changes in CBF, CBV and BOLD that peak some 5 s after the stimulus. We assume that signal changes between activated and baseline states occur instantaneously. Consequently, we can compute the expected signal change for a single pair of activation and control measurements without determining explicitly the block durations that might be required in practice, thus circumventing the need to consider transients. Ignoring transients, we can estimate the time-averaged *CNR*_f_ for an arbitrary number of measurement pairs by multiplying by the square root of the number of averages permissible in the duration of the experiment.

## Results

We present results corresponding to two extreme situations: when a change in CBV is offset entirely by a change in GM tissue volume, *CNR*_f_(*Δ*GM, 30 min), or when that change is offset entirely by a change in CSF volume, *CNR*_f_(*Δ*CSF, 30 min). The actual situation is likely to vary between these extremes depending on the region of brain being activated and the size of the ROI, and we do not know definitively the situation that exists in PMC. While we simulated three sequences for the two conditions—a single polarizing pulse, IR and DIR—the results for the IR and DIR variants did not differ from the results for a single polarizing pulse and there was no advantage in nulling one or more brain constituents (see Supplementary material).

Our result for *CNR*_f_(*Δ*GM, 30 min) as a function of the polarizing time is shown in Fig. 3A, with a maximum of 3.1 at *T_p_* = 840 ms and *T_w_* = 100 ms. The maximum at *T_w_* = 100 ms arises because the GM signal decays twice as rapidly as blood signal at ULF, so that the change in signal is dependent on the difference of the two exponential decays.

**Fig. 3.**
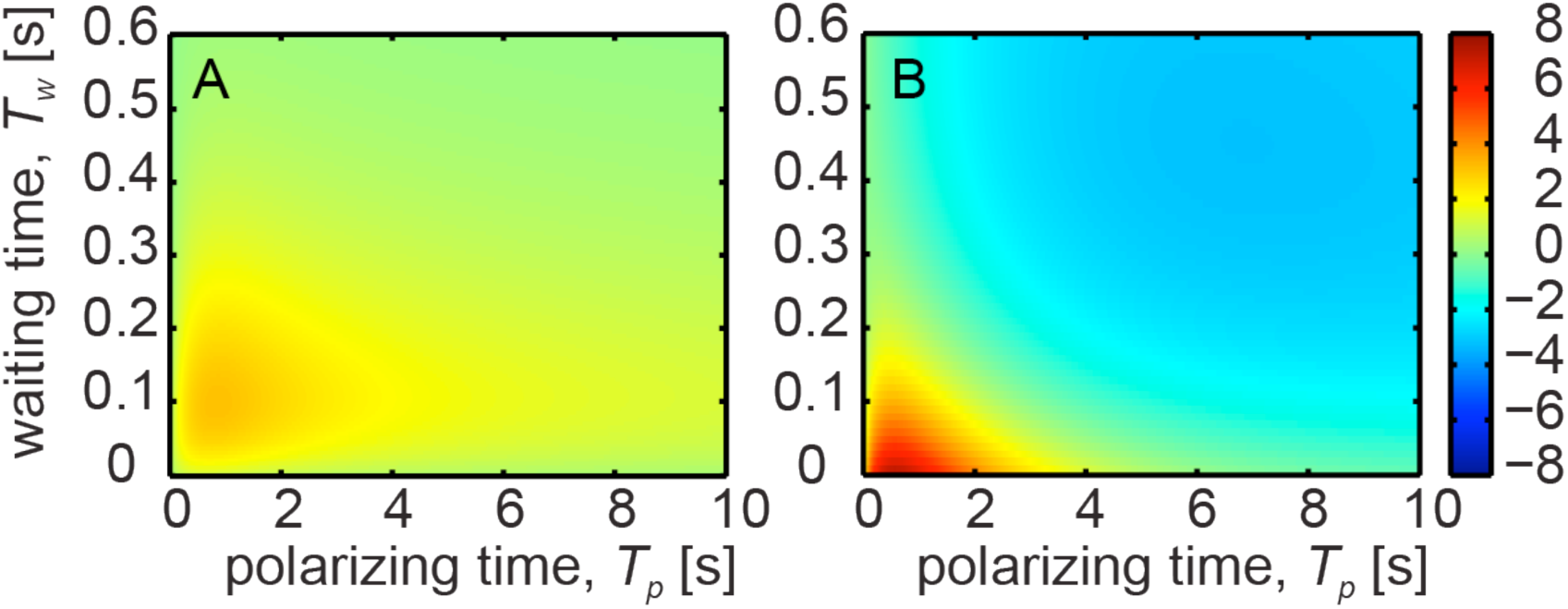
Functional contrast-to-noise ratios. (A) *CNR*_f_(ΔGM, 30 min) as a function of polarizing time *T_p_* and waiting time *T_w_* for a single prepolarizing pulse sequence, for a 10% increase in CBV accommodated by a decrease in GM tissue volume. (B) *CNR*_f_(ΔCSF, 30 min) as a function of polarizing time *T_p_* and waiting time *T_w_* for a single prepolarizing pulse sequence, for a 10% increase in resting CBV accommodated by a decrease in CSF volume.

The maximum *CNR*_f_ is enhanced if the increase in CBV is offset by a decrease in the volume of CSF instead of GM [see Fig. 3B]. The *CNR*_f_(ΔCSF, 30 min) is 7.1 and requires a polarizing time of only 580 ms, with no additional evolution under *B*_0_ beyond the minimum required *T_w_* = 10 ms. The enhanced *CNR*_f_ comes about primarily because there is a larger difference in *T*_1_ at 160 mT between CSF and blood than between GM and blood, together with a small gain in averaging efficiency from the shorter polarizing time. Additional evolution under *B*_0_ beyond *T_w_* = 10 ms produces rapid decay of *CNR*_f_(*Δ*CSF, 30 min), eventually causing a contrast inversion [Fig. 3(B) blue area]. The inversion arises because the long *T*_1_ of CSF creates a persistent signal that is subtracted from a fast-decaying blood signal.

## Discussion

Our simulations for an idealized fMRI experiment at ULF demonstrate that the maximum achievable functional SNR lies between approximately 3 and 7 for a 30-minute averaging time, when using endogenous *T*_1_-based signal changes arising from activation-induced alterations in CBV. In the absence of prior knowledge of the CSF, CBV and GM fractions in the ROI, one needs to choose a timing compromise that might work in practice. The strong dependence on *T*_w_ and *T*_p_ for displacement of CSF [Fig. 3B] but a relatively slow dependence of *T*_w_ and *T*_p_ for displacement of GM [bottom-left corner of Fig. 3A] suggests a compromise setting of *T*_w_ of 50-80 ms and *T*_p_ ~800 ms, for an *CNR*_f_ of between 3 and 5. Our estimated sensitivity indicates that fMRI at ULF is theoretically feasible but challenging, and there are numerous practical matters we must consider to be able to achieve the theoretical sensitivity.

We believe the upgraded technical specifications of the ULFMRI scanner are feasible. Further gains may be made in signal-detection pulse sequences. We assumed a single spin echo acquired for 40 ms, regardless of the *T*_2_-values of signals being detected. For brain tissue, which has *T*_2_ ~ *T*_1_ ~ 85 ms, we might reasonably expect to acquire as many as four echoes although only the first two would have high SNR. Shortening *TE* is non-trivial because of the long *B*_1_ pulses (in the range of 5 ms) and concomitant gradients (Myers et al., 2005). When detecting CSF and blood, however, where *T*_2_ >> 40 ms, we would likely gain significantly by detecting multiple spin echoes for each polarizing period. We did not attempt to optimize the number of spin echoes during the detection period because it complicates the overall sequence timing.

The biggest challenge to setting up an fMRI experiment on a human subject at ULF is probably the temporal stability necessary to realize the *CNR*_f_ we have simulated. Since it is difficult for subjects to sit motionless for 30 minutes in our current ULFMRI system, we could redesign it for the subject to lie supine, as for conventional MRI. Systems for measuring MEG from supine subjects are well established, and it would be possible to adapt existing dewar designs (www.elekta.com). For seated patients, using shorter duration experiments to reduce system and subject instabilities, we predict *CNR*_f_ of between 1.5 and 3.5 for a blocked motor task fMRI experiment in a reduced averaging time of 7.5 minutes.

We emphasize that some of the physiologic assumptions we used may result in differences in the maximum possible *CNR*_f_. The value for blood *T*_1_ at 80 mT was obtained from the arm of a volunteer (Inglis et al., 2013) and blood flow is likely to have contributed some error in that measurement. A 10% error in the low field *T*_1_ of blood used in our simulations changes *CNR*_f_ by only 1%, but a 10% error in high field *T*_1_ changes *CNR*_f_ by 8%. It should be noted that blood *T*_1_ changes with hematocrit and therefore varies across a population (Silvennoinen et al., 2003). It is also somewhat difficult to measure accurately very long *T*_1_ species at ULF, particularly CSF (Inglis et al., 2013). Fortunately, the errors in *T*_1_ for CSF propagate in a relatively benign fashion. An error of 10% in the high field value of *T*_1_ we used for CSF would give roughly a 2% change in *CNR*_f_, while a 10% error in the low field *T*_1_ value alters *CNR*_f_ by less than 1%. Errors in our assumed tissue, CSF and blood fractions in the ROI, as well as their spin densities, propagate linearly in the model.

We ignored the temporal characteristics of CBV changes in our simulations, assuming instantaneous changes of CBV in response to activation. Even though we determined pairwise *CNR*_f_ for computational convenience, neglecting transients is equivalent to assuming the use of a long duration blocked design experiment so that the onset and offset become negligible— that is, the volume fractions for rest and activation blocks are reasonably approximated as steady states. This should be a realistic approximation for blocks lasting tens of seconds, when the transients account for a small fraction of the overall functional signal changes and we can consider the onset and offset delays as a simple phase shift. A long duration blocked design experiment would require good temporal stability for the ULFMRI system, however, as well as minimal head motion.

We opted to ignore BOLD signal changes at ULF, an assumption that is likely to be valid in a real fMRI experiment at *B*_0_ = 130 μT. Other issues we neglected, however, would be important to consider in a real system. We assumed all blood destined for the imaging ROI undergoes the same spin history, and that the brain tissue compartments are in a steady state even though water may exchange between them. Inflow effects from non-uniform preparation of blood outside the head, arising from the heterogeneity of *B_p_*, as well as perfusion-like signal changes arising from the exchange of water from blood to tissue, may alter the magnitude of the fMRI signal change (Lu et al., 2013). The *T*_1_ for blood is rather short at ULF, however, and we expect perfusion effects to be small.

When we began this work, we had assumed that the spread of *T*_1_-values at ULF would provide a significant advantage for CBV-based functional contrast compared to similar methods implemented at high field. In fact, our simulations of IR and DIR sequences converged with the results of a single prepolarizing pulse sequence. Thus, according to our model, the IR and DIR sequences offer no advantage for fMRI at ULF using prepolarization. Instead, the sensitivity of the experiment is primarily determined by the speed with which new signal can be created, that is, by the *T*_1_ of species at *B_p_,* rather than by evolution of the larger *T*_1_ differences at ULF. Small secondary gains are feasible by tuning evolution delays under ULF, as demonstrated by the maximum in Fig. 3A, but our observation that *T*_1_ under *B_p_* drives *CNR*_f_ emphasizes that fMRI at ULF is fundamentally limited by SNR (and instrument stability) rather than by a dearth of intrinsic contrast.

One potential way to enhance the functional changes beyond our simulated limits is judicious use of baseline conditions, for example deactivated states rather than rest. While we contemplated right-versus left-hand finger tapping as a model to activate PMC, we assumed that the ipsilateral cortex would maintain its resting CBV during control blocks rather than having decreased CBV caused by inhibition, yielding a conservative estimate of blood volume change. However, comparing blocks of right-versus left-hand finger tapping reduces BOLD signal in ipsilateral cortex by around 15% (Stefanovic et al., 2004). If we assume a corresponding percentage reduction in the baseline CBV we could expect a commensurate gain in *CNR*_f_. On the other hand, primary cortical areas exhibit stronger BOLD contrast than many brain regions, and as such we should consider the signal changes estimated for PMC to be upper bounds on what might be detected in a brain.

A more radical alternative might be to use exogenous *T*_1_ contrast, as for the first demonstration of functional MRI (Belliveau et al., 1991). An intravascular relaxation agent such as a gadolinium chelate can be expected to reduce the *T*_1_ and *T*_2_ of blood significantly by dipolar relaxation at ULF, as at high field. We simulated a hypothetical situation with blood *T*_1_ shortened to the point that blood signal would be undetectable at *T_w_* = 10 ms and *TE* = 20 ms (see Supplementary material). Fully eliminating blood in this manner produced maximum *CNR*_f_(*Δ*GM, 30 min) and *CNR*_f_(*Δ*CSF, 30 min) of −7.8 and −4.7, respectively, compared to 3.1 and 7.1 using endogenous contrast. These results do not justify moving away from endogenous contrast mechanisms.

Interleaving measurements of functional ULFMRI with MEG measurements in a single device would reduce the duty cycle of MRI sampling, thereby reducing the *CNR*_f_ per unit time. The actual cost would depend upon the system specifications, most critically how quickly it might be switched between MEG and ULFMRI modes. If we assume an MEG sampling period of 500 ms and transitions of 250 ms for MEG to ULFMRI and vice versa, the efficiency of signal averaging for ULF-fMRI decreases by more than 50%. Such a penalty suggests that, at least initially, it would be prudent to aim for longer duration blocks of sequential MEG and ULF-fMRI measurements.

## Conclusions

Our simulations indicate that functional MRI at ULF is feasible provided a sufficiently sensitive ULFMRI magnetic field detector can be implemented. The broad range of *T*_1_ values for brain constituents at ULF offers a viable functional contrast mechanism that can be exploited if the SNR can be made high enough. Measuring the functional signal change we have simulated would then be contingent on signal stability, most critically, as for high field fMRI, on low subject motion.

## Acknowledgements

K.B. thanks the Deutsche Forschungsgemeinschaft for a research fellowship. A.F.P. thanks Oticon for a Master Thesis grant. This work was supported by the Henry H. Wheeler, Jr. Brain Imaging Center at UC Berkeley and the Donaldson Trust.

Author contributions
K.B., A.P., P.S.G., J.C. and B.I. designed research; K.B. and A.P. performed simulations; K.S. verified simulations code; K.B., A.P., J.C. and B.I. analyzed results; and K.B., K.S., J.C. and B.I. wrote the paper.

